# Contribution of arm movements to balance recovery after tripping in older adults

**DOI:** 10.1101/2021.08.03.454896

**Authors:** Sjoerd M. Bruijn, Lizeth H. Sloot, Idsart Kingma, Mirjam Pijnappels

## Abstract

Falls are common in daily life, often caused by trips and slips and, particularly in older adults, with serious consequences. Although arm movements play an important role in balance control, there is limited research into the role of arm movements during balance recovery after tripping in older adults. We investigated how older adults use their arms to recover from a trip and the difference in the effects of arm movements between fallers (n=5) and non-fallers (n=11).

Sixteen older males and females (69.7±2.3 years) walked along a walkway and were occasionally tripped over suddenly appearing obstacles. We analysed the first trip using a biomechanical model based on full-body kinematics and force-plate data to calculate whole body orientation during the trip and recovery phase. With this model, we simulated the effects of arm movements at foot-obstacle impact and during trip recovery on body orientation.

Apart from an increase in sagittal plane forward body rotation at touchdown in fallers, we found no significant differences between fallers and non-fallers in the effects of arm movements on trip recovery. Like earlier studies in young adults, we found that arm movements during the recovery phase had most favourable effects in the transverse plane: by delaying the transfer of angular momentum of the arms to the body, older adults rotated the tripped side more forward thereby allowing for a larger recovery step. Older adults that are prone to falling might improve their balance recovery after tripping by learning to prolong ongoing arm movements.

## Introduction

Falls are common in daily life, particularly in older adults, and a large proportion of these falls are caused by trips and slips (Talbot et al., 2005). The often-seen flailing of the arms after a perturbation makes one think that humans use their arms for balance recovery. Previous work has shown that arm swing during normal walking decreases stability (Bruijn et al., 2010; Meyns et al., 2013; Pijnappels et al., 2010), whereas prolonging ongoing arm swing after a perturbation may be beneficial for balance recovery (Pijnappels et al., 2010).

After tripping, roughly two recovery strategies can be observed; the elevating strategy, in which the tripped foot is placed over the obstacle, and the lowering strategy, in which the tripped foot is placed back on the ground before the obstacle (Eng et al., 1994). Pijnappels et al. (2010) showed that during the elevating strategy, young adults use their arms to change their body orientation mainly in the transverse plane. Prolongation of the ongoing arm movements after obstacle impact by exaggerated shoulder flexion on the tripped side, and exaggerated shoulder extension on the contralateral side, delays the transfer of angular momentum from the arms to the body. This leads to a more favourable body orientation in the transverse plane with the tripped side being rotated more forward, which allows for a larger recovery step.

Although the role of the arms in recovering from a perturbation has mostly been studied in young adults, this role may change with increasing age(Akinlosotu et al., 2020; Merrill et al., 2017; Roos et al., 2008). It has been suggested that, after tripping over an obstacle, older adults exhibit a more protective arm strategy, by reaching both arms forward in anticipation of impacting the floor, rather than using the arms to prevent themselves from falling (Roos et al., 2008). Possibly, older adults who fall after a trip, do so (partly) because of these less effective arm movements for trip recovery.

Thus, we investigated whether and how older adults use their arms to recover from a trip with an elevation strategy. Specifically, we evaluated the difference in the effects of arm movements on body orientation when recovering from a trip between older adults who fell and those who did not fall.

## Methods

### Participants

Sixteen older adults (11 males, mean age 69.7 (SD 2.3) years, mean weight 77 (SD 13)kg, mean height 1.68 (SD 0.09)m) participated. All participants were fit and had no orthopedic, neuromuscular, cardiac or visual problems. All participants signed informed consent, and the protocol was approved by the local ethical committee (#2010-10).

### Procedure

First, participants were fitted with clusters of 3 infrared LED’s for movement registration on the feet, shanks, thighs, pelvis, trunk, upper and lower arms. Kinematics were sampled at 50 samples/second (Optotrak, Northern Digital, Waterloo, Ontario, Canada). Ground reaction forces were sampled at 1000 samples/second. Participants walked repeatedly at a self-selected speed along a 12 by 2.5 m walkway, in which 21 obstacles were hidden. (Pijnappels et al., 2010). Each subject walked on average 64 (SD 10) times along the walkway, in which they were randomly perturbed 7 (SD 2) times on the right leg. Participants were encouraged to take breaks when needed. On average, the first trip occurred after 9 walks.

### Calculations

For this study, we analysed only the first trip, as subsequent trips may contain habituation effects (Akinlosotu et al., 2020; Pijnappels et al., 2010). Tripping responses were classified by visual inspection of 3D kinematics by two independent observers (LHS & SMB) into 1) lowering strategy, 2) successful elevating strategy, 3) unsuccessful elevating strategy. In the current study, we analysed the latter two, and discriminated between participants that fell (‘fallers’, n=5) and those that did not (‘non-fallers’, n=11).

We calculated angular momenta (*L)* of all segments around the total body center of mass (CoM), as well as the combined inertia of trunk and legs (*I*_*trunklegs*_) with respect to the CoM (Pijnappels et al., 2010). We estimated the isolated effects of arm movements at foot-obstacle impact (“impact”) and during the recovery phase, lasting from impact until touchdown of the recovery foot (“touchdown”)(Pijnappels et al., 2010). First, we calculated the body orientation during the recovery phase (Actual). Second, we calculated how the body would have rotated if the angular momentum of the arms at impact would be instantaneously transferred to the rest of the body, without allowing any further arm movements after impact (Transfer & Cut). Third, we calculated how the body would have rotated if no angular momentum of the arms would be transferred to the body at impact, and the arms would prolong their movements throughout the recovery phase (Cut). For each of these three angular momenta curves (*L*_*calculation*_), we calculated the angular velocity (*ω*_*calculation*_) from:

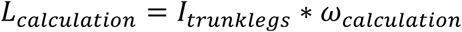

This angular velocity was subsequently integrated over the recovery phase, starting from 0^0^ at impact, to obtain the 3D orientation of the body at touchdown (figure 1). For mathematical details, see (Pijnappels et al., 2010).

**Figure 1:**
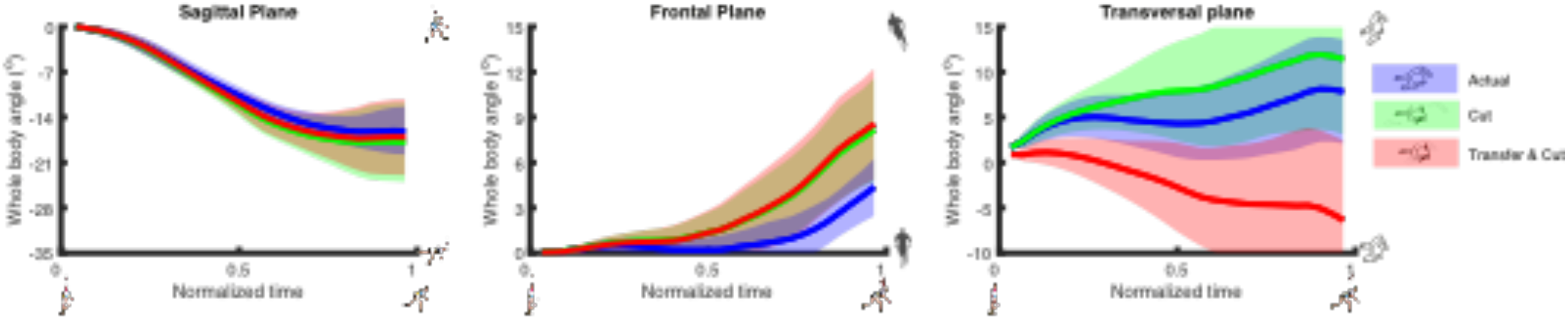
The effects of Calculation mode (different colours) on the total body orientation (y-axis) as a function of normalised time (x-axis, 0 = impact, 1.0= touchdown) in the Sagittal (left), Frontal (middle), and Transverse (right) planes. Data represent the mean for the non-fallers. For each panel, the figures on the right illustrate what each orientation indicates. Shaded regions represent standard deviations.

### Statistical analysis

We compared pre-tripping walking speed and time between impact and touchdown between fallers and non-fallers using an unpaired t-test. To test for differences in the effects of arm movements on body orientation at touchdown for each plane, we used a mixed model ANOVA, with Group (faller, non-faller) as between factor, and Calculation mode (Actual, Transfer & Cut, Cut) as within factor. Significant effects were followed up by paired t-tests with Bonferroni correction. All analyses were performed in Matlab (R2019A, Nattick, Massachusetts: The MathWorks Inc.), with *α*=0.05.

## Results

There were no temporal differences in recovery between fallers and non-fallers. Walking speed of the fallers (1.48m/s (SD 0.21)) was not significantly different from the non-fallers (1.43m/s (SD 0.07), p=0.48). Also, time between impact and touchdown of fallers (464ms (SD 84)) did not significantly differ from non-fallers (496ms (SD 61), p=0.40). Thus, there were no differences in the time over which angular velocities were integrated.

Arm movements had an effect on trip recovery in all three planes, and this effect was not different between fallers and non-fallers. In the sagittal plane, the Cut calculation led to small, although significantly more forward body rotation of about 1° than the Transfer & Cut calculation (figures 1 & 2, table 1 & 2). Thus, in the sagittal plane, it may be undesirable to delay transfer of angular momentum from the arms to the body, as this would lead to more sagittal plane forward body rotation (as in the Cut calculation). In the frontal plane, the Actual calculation led to a significantly more vertical body orientation than both the Cut and Transfer & Cut calculations, which were more laterally rotated towards the tripped side. This indicates that participants were able to fully cancel all angular momentum that was present at impact (i.e. the Transfer & Cut calculation), and even could cancel angular momentum from the body (since the actual orientation of the body was rotated significantly less than the Cut calculation). Effects of arm movements were largest in the transverse plane. In this plane, the Actual calculation led to an orientation with the tripped side rotated significantly less forward than the Cut calculation, yet rotated significantly more forward than the Transfer & Cut calculation. This indicates that in the transverse plane, participants benefitted from delaying transfer of angular momentum from the arms to their body, so much even, that would they not do so, the tripped right side of their body would have been rotated backward at recovery foot touchdown. Still, participants did not manage to delay transfer of all arm angular momentum, as indicated by a significant difference between Actual and Cut calculations.

**Table 1:** Results of the statistical tests. Significant effects are displayed in bold.

|  | Calculation mode |  | Fallstatus |  | Calculation mode X Fallstatus |  |
| --- | --- | --- | --- | --- | --- | --- |
|  | F(2,28) | p | F(F1,14) | p | F(2,28) | p |
| Sagittal | 5.04 | 0.014 | 9.82 | 0.007 | 0.85 | 0.438 |
| Frontal | 16.74 | <0.001 | 0.05 | 0.831 | 1.18 | 0.324 |
| Transverse | 67.34 | <0.001 | 0.46 | 0.508 | 0.70 | 0.507 |

**Table 2:**
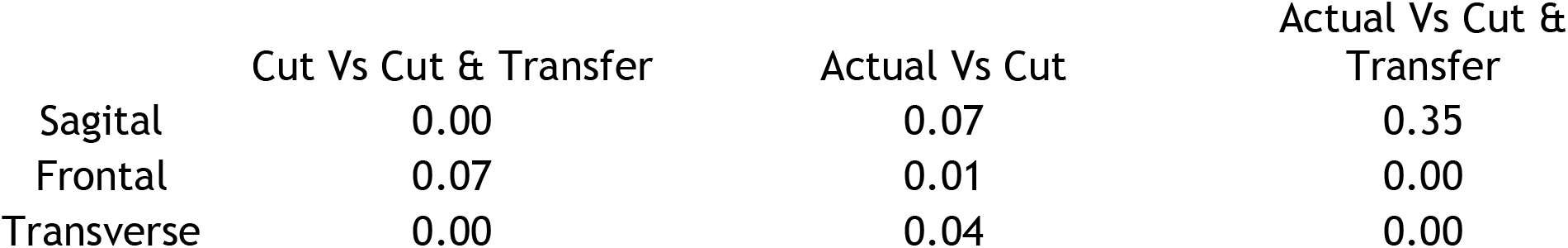
Posthoc results. Significant effects are displayed in bold.

There were only significant differences between fallers and non-fallers in the sagittal plane; Fallers had a more Actual sagittal plane forward body rotation at touchdown than non-fallers (figure 2, table 1).

**Figure 2.**
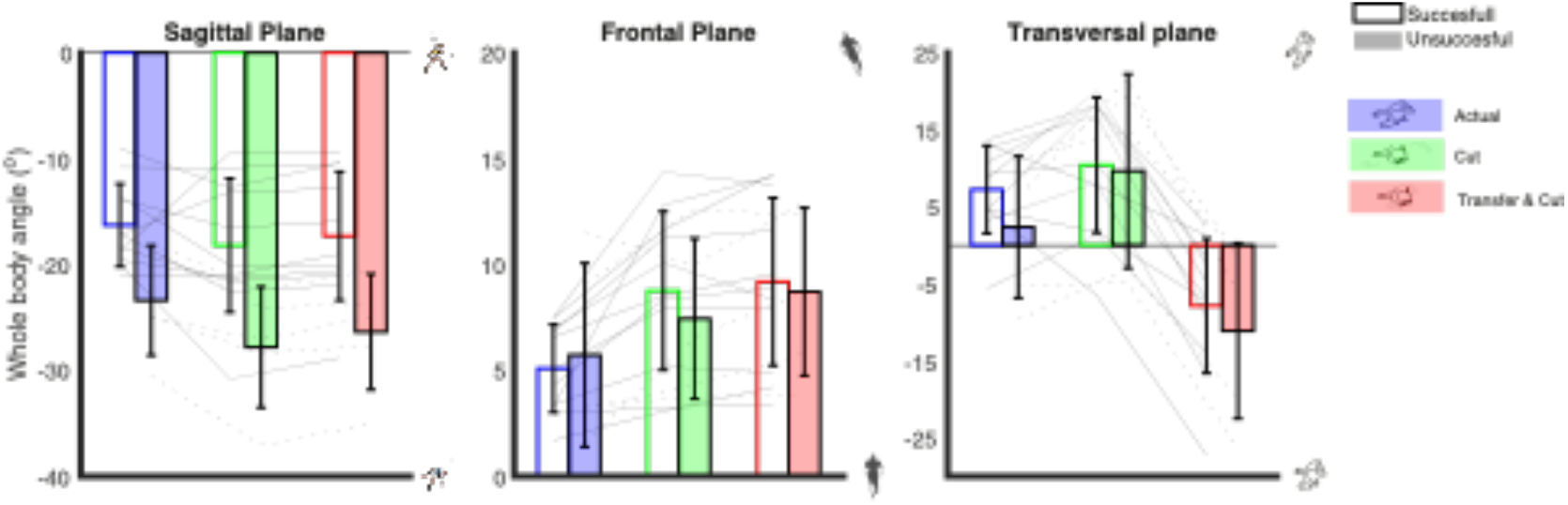
Body orientation at touchdown for non-fallers (non-filled bars) and fallers (solid colours) as a function of Calculation mode (colours). Error bars represent standard deviations, and lines represent individual data.

## Discussion

We studied the effects of arm movements on recovery after a trip in older fallers and non-fallers. Similar to earlier studies in young adults, we found that arm movements had most effect in the transverse plane, where older adults (regardless of fall-status) delayed the transfer of angular momentum of the arms to the body (i.e. behaved like the Cut calculation). This allowed them to gain a more favourable body orientation in the transverse plane, with the tripped right side rotated more forward, thereby lengthening the recovery step. Apart from an increase in sagittal plane forward body rotation in fallers compared to non-fallers, we found no significant differences between fallers and non-fallers.

Interestingly, the effects of arm movements did not significantly differ between fallers and non-fallers in the transverse plane. A post-hoc analysis showed that fallers (0.56 (SD 0.14)m) took a significantly shorter recovery step than non-fallers (0.82 (SD 0.11)m, P<0.01). Thus, falling after a trip seems to be related to problems with step lengthening, which in our sample could not be connected to ineffective arm movements. Although Roos et al. (2008) suggested that older adults use a protective strategy of extending both arms to negotiate impact in case of a fall, which would be destabilizing according to our analyses, we did not observe such strategy in our older participants. Alternatively, earlier work suggests that problems in generating hip, knee and ankle moments during push-off may be a more crucial in successful recovery from a trip than the contribution of arm movements (Pijnappels et al., 2005).

Another approach to understanding the importance of arm movements during trip recovery in older adults is to compare our results to findings in young adults. In the sagittal plane, body orientation at touchdown in our older non-fallers seems more vertical than in young adults (−20° forward rotation in (Pijnappels et al., 2010), ∼16° in our study). While this could indicate that older non-fallers were better at recovering from a trip, it could also be related to the fact that they generally walked slower, thus carrying less momentum (1.54m/s in (Pijnappels et al., 2010), 1.48m/s in ours). Moreover, other factors, like push-off ability mentioned previously could be underlying differences in orientation at touchdown.

In the frontal plane, our older adults’ Actual body orientation at touchdown was comparable to that of young adults (Pijnappels et al. (2010), even though the effect of arm movements differed. In the older adults, both the Transfer & Cut and Cut calculations led to significantly more lateral rotation towards the tripped side. Thus, our results suggest that in the frontal plane, older adults were able to cancel the angular momentum of the arms at impact, and also to transfer some of the (hindering) angular momentum from the body to the arms. How they were able to do so is an interesting topic for further investigation.

In the transverse plane, older adults rotated their tripped side forward by only 6.5° versus 18° reported for young adults (Pijnappels et al. (2010), which indicates a substantially more favourable body orientation for the latter group. Like in young adults, arm movements in older adults contributed to this orientation at touchdown. However, there was also a slight but significant difference between the Actual and Cut calculations, implying that older adults did not cancel all angular momentum of the arms at impact. Thus, older adults could perhaps improve their recovery by further prolonging the ongoing movements of the arms, thereby delaying the transfer of angular momentum from the arms to the body, achieving more forward rotation of the tripped side, like younger adults, and potentially increasing recovery step length.

### Limitations

Our number of participants, particularly fallers, was small, so we may have missed smaller effects. Second, kinematics data was sampled at a relatively low sample rate of 50Hz. However, the analyzed trip recovery period was 400ms, and arm and trunk motions are unlikely to show substantial motions at >10Hz.

## Conclusion

Arm movements during tripping in older adults help to move the body in a more favourable orientation for balance recovery in the transverse plane. While the recovery step was smaller in fallers, we found only minor differences in body orientation and arm contribution between fallers and non-fallers. This suggests that arm movements are not a major factor differentiating fallers from non-fallers.

## Acknowledgements

SMB was funded by a VIDI grant (016.Vidi.178.014) from the Dutch Organization for Scientific Research (NWO), SMB and MP were funded by a by a VIDI grant (VIDI grant (no. 91714344)) from the Dutch Organization for Scientific Research (NWO)

## Notes

### Competing Interest Statement

The authors have declared no competing interest.

### Summary of Updates

Revision based on peer-review suggestions from Journal of Biomechanics., minor revisions, round 2

